# Connectivity and functional diversity of different temporo-occipital nodes for action perception

**DOI:** 10.1101/2024.01.12.574860

**Authors:** Baichen Li, Marta Poyo Solanas, Giuseppe Marrazzo, Beatrice de Gelder

## Abstract

The temporo-occipital cortex (TOC) plays a key role in body and action perception, but current understanding of its functions is still limited. TOC body regions are heterogeneous and their role in action perception is poorly understood. This study adopted data-driven approaches to region selectivity and investigated the connectivity of TOC nodes and the functional network sensitivity for different whole body action videos. In two human 7T fMRI experiments using independent component analysis, four adjacent body selective nodes were detected within the TOC network with distinct connectivity profiles and functional roles. Action type independent connectivity was observed for the posterior-ventral node to the visual cortex, the posterior-dorsal node to the precuneus and the anterior nodes to the frontal cortex. Action specific connectivity modulations were found in middle frontal gyrus for the aggressive condition with increased connectivity to the anterior node and decreased connectivity to the posterior-dorsal node. But for the defensive condition, node-nonspecific enhancement was found for the TOC-cingulate connectivity. By addressing the issue of multiple nodes in the temporo-occipital network we show a functional dissociation of different body selective centres related to the action type and a potential hierarchy within the TOC body network.

## 1. Introduction

During social interactions, intentions, actions, and emotions are routinely read from nonverbal communication signals provided by faces, body postures and whole-body movements as well-documented in studies of humans (Argyle, 1976; de Gelder et al., 2010) and non-human primates (Vogels, 2022). Despite its importance, the neural basis of whole-body perception is still much less understood than that of faces (Deen et al., 2023).

Recent studies have revealed several brain areas in human and non-human primates. In humans, the extrastriate body area (EBA) (Downing & Kanwisher, 2001) was the first one to be reported. It is a region located in the extrastriate cortex that overlaps with other category-selective areas, such as those dedicated to processing motion (Weiner & Grill-Spector, 2011), tools, and even action related words (Lingnau & Downing, 2015). Subsequent studies have identified at least three different body-selective clusters within the extrastriate cortex (Weiner & Grill-Spector, 2011). This anatomical diversity is complemented by functional diversity, as these clusters also display varying patterns of connectivity (Zimmermann et al., 2018).

The actual contribution of these body selective areas is not well understood (de Gelder & Poyo Solanas, 2021; de Gelder et al., 2010; Vogels, 2022). A number of studies found that these body selective areas play a role in processing emotional expressions (de Gelder et al., 2004; Grèzes et al., 2007; Peelen et al., 2007; Pichon et al., 2008), biological movement (Jastorff & Orban, 2009), specific postural and kinematic features of the body (Marrazzo et al., 2023; Marrazzo et al., 2021; Poyo Solanas et al., 2020), action recognition (Goldberg et al., 2014; Shmuelof & Zohary, 2005), motor planning (Zimmermann et al., 2012), as well as social perception (Kret et al., 2011a; Moreau et al., 2023). Taken together, these findings suggest that body selectivity may be understood not simply as a matter of category selectivity but as resulting from the activity of a sparsely distributed ensemble of brain areas (de Gelder & Poyo Solanas, 2021; Weiner & Grill-Spector, 2011). Mapping the broader network’s activity may be an important step in uncovering how these different body-specific nodes collectively contribute to the perception of whole-body movements at the network level. Several network models have been proposed including the Action Observation Network (AON) (Caspers et al., 2010), the Default Mode Network (DMN) (Zhan et al., 2018) or a pathway for social perception (Haak & Beckmann, 2018). However, none of these directly addresses the specific case of perceiving emotional expressions, intentions, and actions of conspecifics routinely conveyed by body movements.

The goal of this study was to identify a dynamic whole-body network and to investigate how its network activity and its connectivity with other brain areas supports specific actions such as defensive or aggressive behaviour. Rather than following the traditional approach of contrast-based selection of body regions, we approach the question at the network level with data-driven methods. We used independent component analysis (ICA), which is widely applied in resting-state and task-based fMRI studies (Du et al., 2017; Jarrahi et al., 2015; Jung et al., 2020), to identify the body sensitive nodes within the TOC network and further tracked their whole-brain communications during whole body action processing.

## 2. Method

The study consisted of two experiments: a localizer experiment and the main experiment. First, we used the localizer experiment to identify the temporo-occipital network associated with body action perception. This was accomplished through a data-driven strategy based on our previous study (Li et al., 2023). Next, the data of the main experiment was employed to extract node regions within this network and investigate their connectivity profiles as well as their modulation by affective body conditions. Nineteen participants took part in the experiment.

### 2.1. Participants

Nineteen healthy participants (mean age = 24.58 years; age range = 19-30 years; 6 males, all right-handed) took part in the experiment. All participants had a normal or correct-ed-to-normal vision and no medical history of any psychiatric or neurological disorders. All participants provided informed written consent before the start of the experiment and received a monetary reward (vouchers) or course credits for their participation. The experiment was approved by the Ethical Committee at Maastricht University and was performed in accordance with the Declaration of Helsinki.

### 2.2. Experiment Design

#### 2.2.1. Network localizer

The functional localizer followed a blocked design with twelve categories of videos com-posed from three factors: (body / face / object) * (human / monkey) * (normal / scramble). Each category consisted of ten different 1000-ms videos which were presented block-wise in a random order. Within each block, the ten videos were interleaved by a fixed 500-ms inter-trial interval, while two consecutive blocks were interleaved by an inter-block interval jittered around 11 seconds. The order of block conditions was randomized for each participant, and each condition was repeated six times within three runs. Each run contained a catch block where, in one of the trials, the fixation point changed its shape from a “+” to a “o”. Participants were instructed to make a button-press response when detecting the fixation shape change. The total length of each run was 735 seconds on average. A detailed description of the localizer stimuli and design can be found in Li et al. (2023).

#### 2.2.2. Main experiment

For the main experiment, we applied a mixed block/event-related design (Visscher et al., 2003) consisting of five conditions of videos: three human body conditions (aggressive, defensive, and neutral), one neutral human face condition and one neutral object condition (**Figure 1**). Each condition consisted of ten different 1000-ms videos. The body and face videos were chosen from the same stimulus set as described in Kret et al. (2011b). The object videos were selected from the same set as used in the localizer experiment but differed from the ones used in that experiment.

**Figure 1.**
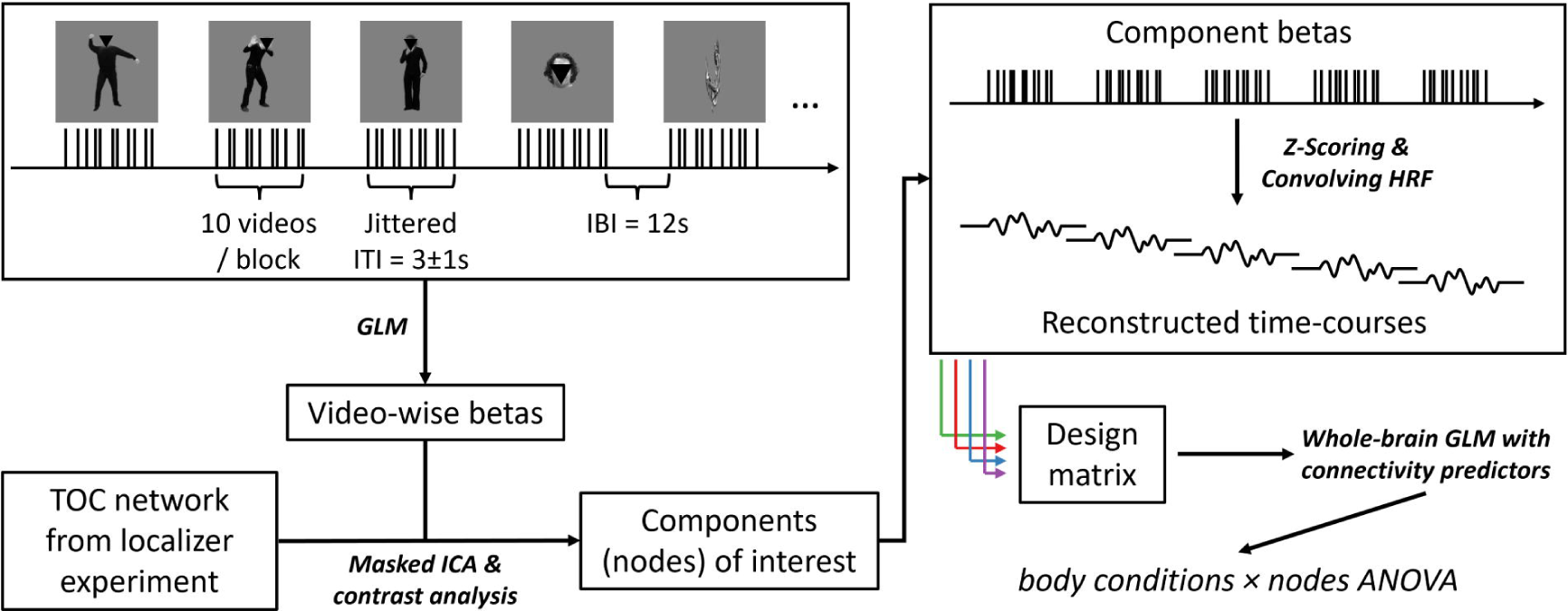
Illustration of the main experiment design and analysis. Five video conditions (aggressive / defensive / neutral face blurred body, neutral face, and object; face region overed here for privacy) were presented in a mixed block/event-related design, in which the stimuli were blocked for each condition while with jittered inter-trial-interval around 3s. For each condition, ten different videos were included and were repeated ten times across five runs. GLM wa conducted to estimate the response for each different video, resulting in 50 betas extracted for each participant. The video-wise betas were then entered to an ICA procedure within the body sensitive TOC network identified by the localizer experiment. Body selective network node were defined by higher component responses for body videos than for non-body videos. To track the whole-brain connectivity of each selected node, the video-wise betas were z-scored and convolved with the hemodynamic response function within each condition, resulting in five reconstructed time-courses for each component. The reconstructed time-courses for all selected components were then added to a whole-brain GLM design matrix as the predictors for seed-based connectivity. Finally, two-factor ANOVA was conducted with connectivity betas across all participants to test their modulations from the body conditions or the node components.

During the experiment, each condition was presented block-wise with a jittered inter-trial interval of around 3 seconds and an inter-block interval of 12 seconds. For each trial, the video was centered and presented on a uniform gray background. The sizes of stimuli were 3.5*7.5 degrees of visual angle for bodies and objects, and 3.5*3.5 degrees of visual angle for faces. The order of block conditions was randomized for each participant, and each condition was repeated ten times within five runs. Two extra blocks with a catch trial were inserted in each run where participants were instructed to detect the fixation shape changes as described in the localizer experiment. The total length of each run was 480 seconds.

Both the main experiment and the localizer experiment were programmed using the Psychtoolbox (https://www.psychtoolbox.net) implemented in Matlab 2018b (https://www.mathworks.com). Stimuli were projected onto a screen at the end of the scanner bore with a Panasonic PT-EZ57OEL projector (screen size = 30 * 18 cm, resolution = 1920 * 1200 pixel). Participants viewed the stimuli through a mirror attached to the head coil (screen-to-eye distance = 99 cm, visual angle = 17.23 * 10.38 degrees).

#### 2.2.3. fMRI data acquisition

All images were acquired with a 7T MAGNETOM scanner at the Maastricht Brain Imaging Centre (MBIC) of Maastricht University, the Netherlands. Functional images were collected using the T2*-weighted multi-band accelerated EPI 2D BOLD sequence (TR/TE = 1000/20 ms, multiband acceleration factor = 3, in-plane isotropic resolution = 1.6 mm, number of slices per volume = 68, matrix size = 128 * 128, volume number = 735 for the network localizer and 480 for the main experiment). T1-weighted anatomical images were obtained using the 3D-MP2RAGE sequence (TR/TE = 5000/2.47 ms, Inverse time TI1/I2 = 900/2750 ms, flip angle FA1/FA2 = 5/3°, in-plane isotropic resolution = 0.7 mm, matrix size = 320 * 320, slice number = 240). Physiological parameters were recorded via pulse oximetry on the index finger of the left hand and with a respiratory belt.

#### 2.2.4. fMRI image preprocessing

Anatomical and functional images were preprocessed using the Brainvoyager 22 (Goebel, 2012), and the Neuroelf toolbox in Matlab (https://neuroelf.net/). For anatomical images, brain extraction was conducted with INV2 images to correct for MP2RAGE background noise. The resolution was then downsampled to 0.8 mm for better alignment to the 1.6 mm resolution of functional images. For functional images, the preprocessing steps included EPI distortion correction (Breman et al., 2020), slice scan time correction, 3D head-motion correction, and high-pass temporal filtering (GLM with Fourier basis set of 3 cycles, including linear trend). Coregistration was first conducted between the anatomical image and its most adjacent functional run using a boundary-based registration (BBR) algorithm (Greve & Fischl, 2009), and all the other functional runs were coregistered to the aligned run. Individual images were normalized to Talairach space (Collins, Neelin, Peters, & Evans, 1994) with 3 mm Gaussian spatial smoothing. Trilinear/sinc interpolation was used in the motion correction step, and sinc interpolation was used in all the other steps.

Physiological parameters were collected as confound factors for the functional imaging data. The physiological data were preprocessed using the RETROspective Image CORrection (RETROICOR; Glover et al., 2000; Harvey et al., 2008) pipeline, which uses Fourier expansions of different orders for the phase of cardiac pulsation (3rd order), respiration (4th order) and cardio-respiratory interaction (1st order). Eighteen physiological confound factors were finally created for each participant.

The anatomical labeling of the brain areas reported in this study was performed according to the Talairach Daemon (http://www.talairach.org/daemon.html) in combination with the Multilevel Human Brain Atlas (https://ebrains.eu/service/human-brain-atlas).

### 2.3. Data analysis

#### 2.3.1. Body network extraction

The Infomax algorithm implemented in the Group ICA of fMRI Toolbox (GIFT; Calhoun et al., 2001) was used to identify body selective networks within the localizer experiment. This resulted in 75 spatial independent components. Individual ICs were back-reconstructed using the GIG-ICA algorithm from the aggregated group ICs (Du & Fan, 2013). The stability of group ICA was assessed by the ICASSO module implemented in the GIFT, which repeated the Infomax decomposition 20 times and resulted in an index of stability (Iq) for each IC (Himberg et al., 2004). Prior to the group-ICA, physiological and motion confounds were regressed out from the preprocessed functional images. The resulting time courses were then transformed into percentages of signal change to enhance the ICA stability (Allen et al., 2011). Components showing large white matter / cerebrospinal fluid coverage were excluded from further analysis.

To identify body selective networks, we conducted a GLM on each reconstructed subject-level IC time course, which estimated the IC response for each condition. In the design matrix, each condition predictor was modeled as a boxcar function with the same duration of the block and convolved with the canonical HRF. The estimated betas were first averaged across all runs for each participant and were then used to calculate the contrast of [2 * human body (normal - scramble) – (human face (normal - scramble) + human object (normal - scramble))]. Right-tailed t-tests and Benjamini-Hochberg multiple comparison corrections were conducted at the group level to find significant body sensitivity.

To define the group-level coverage of the IC networks, the individual IC maps were normalized to z-scores and averaged across all runs for each participant. A group t-test against zero was computed using the z-scored maps of each subject and corrected using a cluster-threshold statistical procedure based on Monte-Carlo simulation (initial p < 0.005, alpha level = 0.05, iteration = 5000). The group-level coverage of the network was then used as the initial mask for the body node extraction in the main experiment.

#### 2.3.2. Connectivity of the TOC body nodes

The analysis for the main experiment is illustrated in **Figure 1**. First, a fixed-effects GLM was conducted on each participant’s functional images with each different video as a separate predictor. Fifty betas were estimated on each voxel for each participant. Next, a group ICA was conducted on the estimated betas within the TOC body network, following its definition with the network localizer. Fifteen ICs were extracted along with their video-wise betas. The component betas were then averaged for each condition and entered a group-level t-test for the contrast of [(aggressive body + defensive body + neutral body) > (face + object)] to select the body nodes.

To track the connectivity between the body nodes and the rest of the brain, a whole brain GLM was conducted with the IC responses as predictors. An event-related design matrix was first constructed by convolving each stimulus duration with the HRF and binned for each condition. Next, since the IC betas were extracted video-wise, the time-courses of each IC could be reconstructed by convolving the betas with a canonical HRF according to the on/offsets of the corresponding videos. These reconstructed time courses were then incorporated into the GLM design matrix to reflect their connectivity. Moreover, the IC betas were normalized within each condition so that the reconstructed time-courses reflected the item-to-item variance while omitting the categorical baseline modulations. The IC time-courses were modeled separately for each IC and each condition. The resulting betas were then entered in a voxel-wise ANOVA with factors Body conditions (aggressive / defensive / neutral) and Seeds nodes to assess the action type modulated connectivity. Statistical maps of significant main and interaction effects were corrected with the Monte Carlo cluster-threshold (initial p < 0.005, alpha level = 0.05, iteration = 3000). Further multiple comparisons and simple effect tests were conducted at the ROI level for each significant cluster.

## 3. Results

Among the nineteen participants, two of them were excluded from both experiments and an extra one was excluded from the main experiment analysis due to large distortion of the functional or anatomical images.

In the localizer experiment, 75 independent components (ICs) were extracted from each subject’s preprocessed functional images. Noise-induced ICs were identified excluded according to the spatial overlap with the white matter / cerebrospinal fluid mask, the mean response, and the r^2^ of the general linear modelling (GLM) fitting on the IC time-course. The details of the criteria are described in Li et al. (2023). By conducting GLM on the IC time-courses, the body-selective occipital network was identified by the analysis of the contrast [2 * human body (normal - scramble) > human face (normal - scramble) + human object (normal - scramble)]. It exhibited a significant preference for human bodies over objects (**Figure 2a**, t(16) = 4.24, Benjamini-Hochberg False Discovery Rate corrected q = 0.006, right-tailed) in the bilateral lateral occipital cortex (LOC) and also included bilateral fusiform cortex, superior parietal lobe (SPL), posterior superior temporal sulcus (pSTS) / temporoparietal junction (TPJ), pulvinar and amygdala.

**Figure 2.**
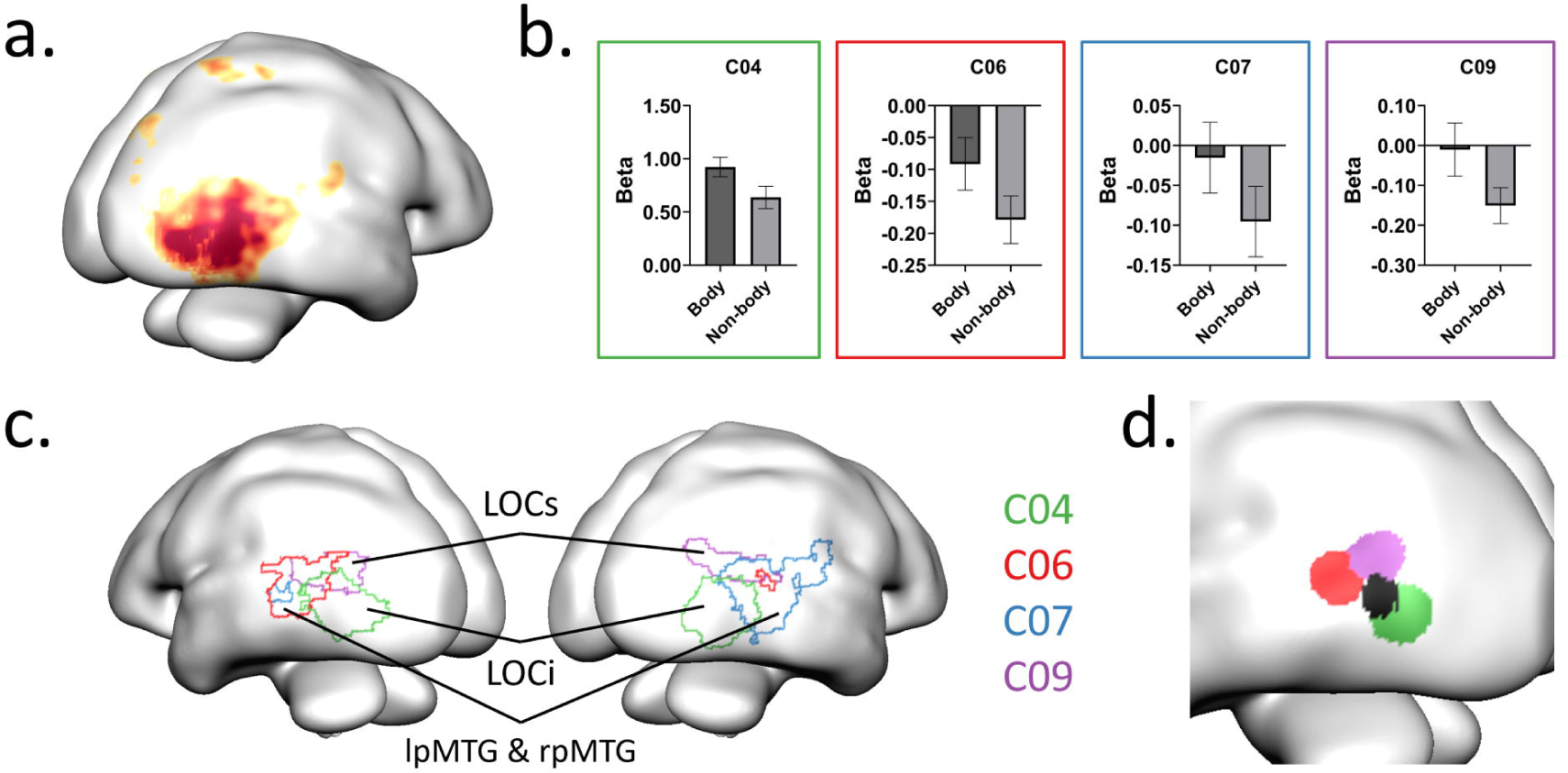
(a). The coverage of TOC network as defined in the network localizer experiment. (b). Beta plots of the four body-nodes from the main experiment. Zero-point indicates the mean beta value across all masked voxels. Colors indicate the component indexes. (c). The spatial distribution of the four body nodes. (d). The relative position of the centers of body nodes and the hMT (black) on the left hemisphere.

Following the identification of body network defined above (referred to as TOC network for simplicity), we examined their connectivity patterns and how they were influenced by the affective conditions. This analysis utilized the data from the main experiment, where 50 betas were extracted for each subject. A GIG-ICA procedure was then conducted within the predefined TOC network on each subject’s 50 betas, resulting in 15 ICs along with their video-wise betas. The component betas were then averaged by condition and entered a group-level t-test for the contrast of [(aggressive body + defensive body + neutral body) > (face + object)] to select body nodes. After multiple comparison correction, four adjacent nodes showed significant body selectivity (C04, t(15) = 8.08, corrected q < 0.001; C06, t(15) = 4.21, corrected q = 0.002; C07, t(15) = 2.66, corrected q = 0.033; C09, t(15) = 4.90, corrected q < 0.001; all right-tailed; **Figure 2b, c**). The decomposed beta values are shown in **Figure 2b**. Since the data was demeaned before entering the ICA, the zero-point in the plots indicate the averaged beta value across all masked voxels. The nodes C04 and C09 were distributed bilaterally covering the inferior LOC (**Figure 2c**, green component, LOCi for abbreviation) and the superior LOC (**Figure 2c**, purple component, LOCs for abbreviation) regions, respectively. The C06 and C07 had a unilateral distribution and covered the posterior middle temporal gyrus on the left (**Figure 2c**, red component, lpMTG) and right (**Figure 2c**, blue component, rpMTG) hemispheres, respectively. Consistent with a previous study on subdivisions of EBA (Weiner & Grill-Spector, 2011), these body nodes partially overlapped with and surrounded the hMT in each hemisphere (**Figure 2d**).

By comparing the spatial maps of the four independent components, we further investigated the voxels which were specifically dominant in each of the nodes. As shown in **Figure 3a**, non-overlapping clusters were identified where the component weights exhibited significantly larger values for one of the components than for the other three. The composition of each cluster was then labelled following the nomenclature of the Human Connectome Project multi-modal parcellation (HCP-MMP; Glasser et al., 2016). This showed that the dominance of the LOCi node was mainly found in V4t, PH, LO2, and FST (**Figure 3b**). The dominance of the pMTG nodes included PHT and TPOJ2 on the left side (**Figure 3b**), and further extended to TPOJ1 on the right side (**Figure 3b**). For the LOCs node, dominance was found especially in TPOJ3 (**Figure 3b**).

**Figure 3.**
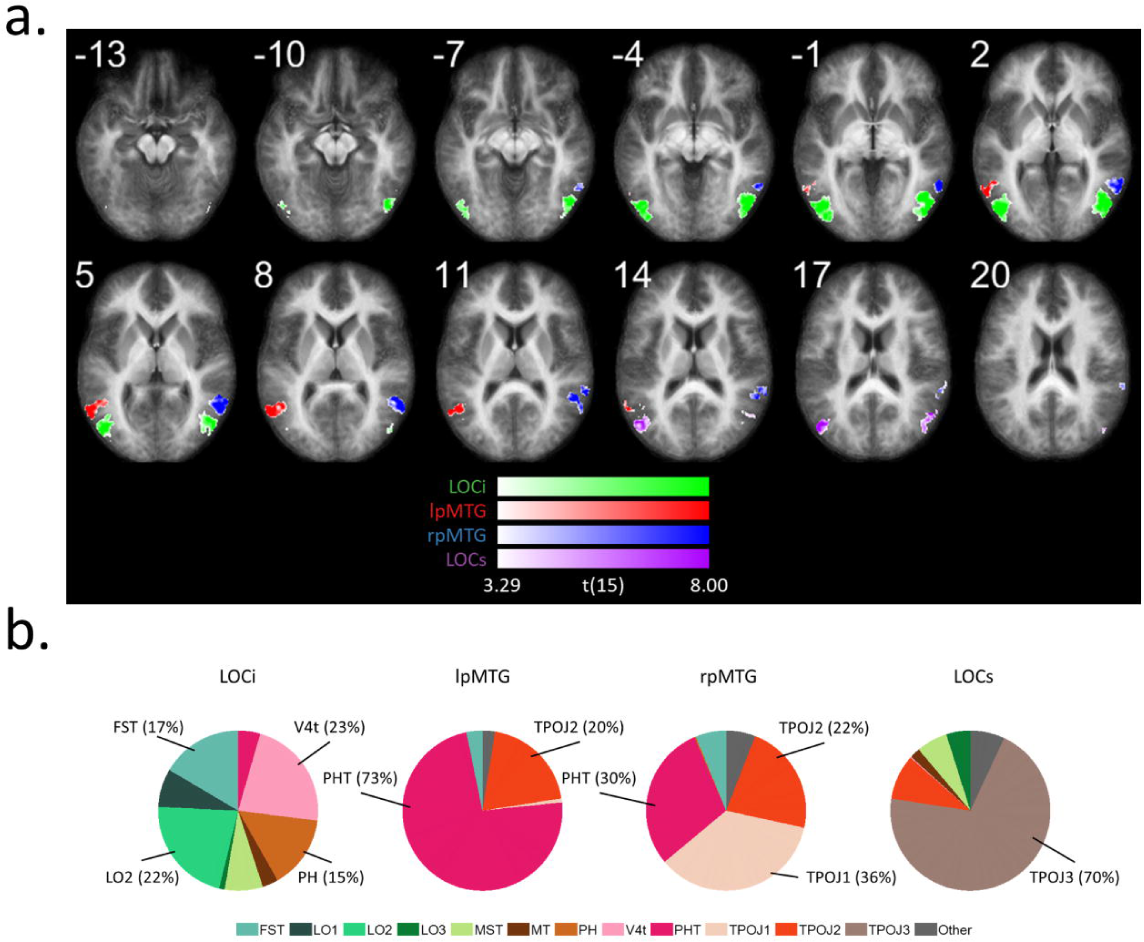
(a). Map of voxels with significantly higher contribution from each of the nodes. The voxel-wise IC weight from each node was compared to the other three nodes and entered a group-level t-test against zero (two-tailed). The resulting map was corrected by a cluster-threshold statistical procedure based on Monte-Carlo simulation (initial p < 0.005, alpha level = 0.05, iteration = 5000). Slice numbers indicate the Z coordinates in the Talairach space. (b). The voxel composition for each cluster in (a.), labeled by the HCP-MMP atlas (Glasser et al., 2016).

To track the connectivity between the body nodes and the rest of the brain, a whole brain GLM was conducted with the IC responses as predictors. Since the IC betas were extracted item-wise, the time-courses of each IC can be reconstructed by convolving the betas with a canonical hemodynamic response function (HRF) according to the on/offsets of the corresponding videos. The IC time-courses were modeled separately for each IC and each condition, resulting in 5 conditions * 4 nodes (LOCi, LOCs, lpMTG, & rpMTG) = 20 connectivity terms added to the GLM. Next, to assess connectivity modulated by the action type, the resulting betas were entered in a voxel-wise ANOVA with the 3 body conditions (aggressive / defensive / neutral) * 4 nodes. A significant main effect of node was found in widely distributed clusters, showing that different TOC body nodes have differentiated connectivity profiles (**Figure 4**; **Table 1**). The LOCi and LOCs nodes showed stronger connectivity to visual cortex and posterior cingulate cortex (PCC) while the two pMTG nodes were connected more dominantly to the middle & posterior insula, supramarginal gyrus (SMG) and frontal regions (**Figure 4**; **Table 1**).

**Figure 4.**
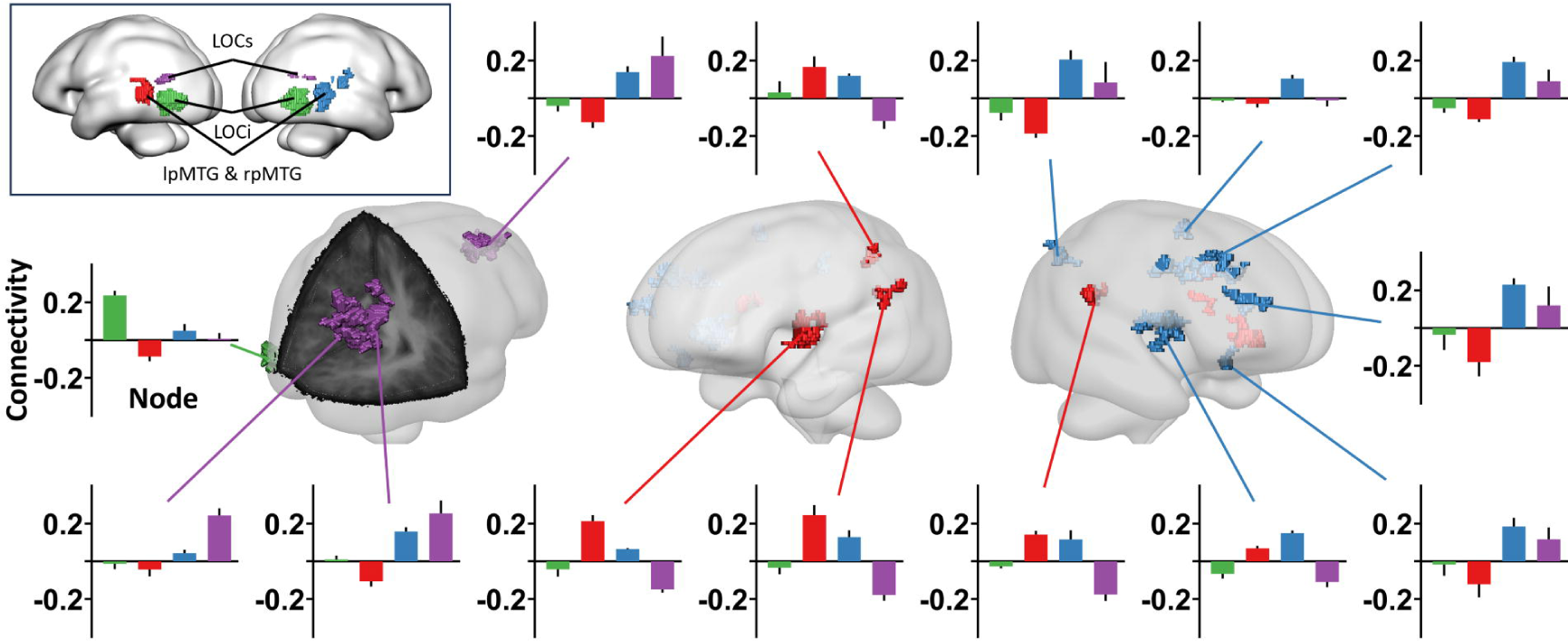
Clusters with significant main effect of the seed node on connectivity. Cluster color indicate the seed nodes with the highest connectivity to the corresponding clusters. Bar plots show the detailed connectivity profile of each cluster, with bar colors indicating the seed nodes as shown in the upper-left framed panel.

**Table 1.**
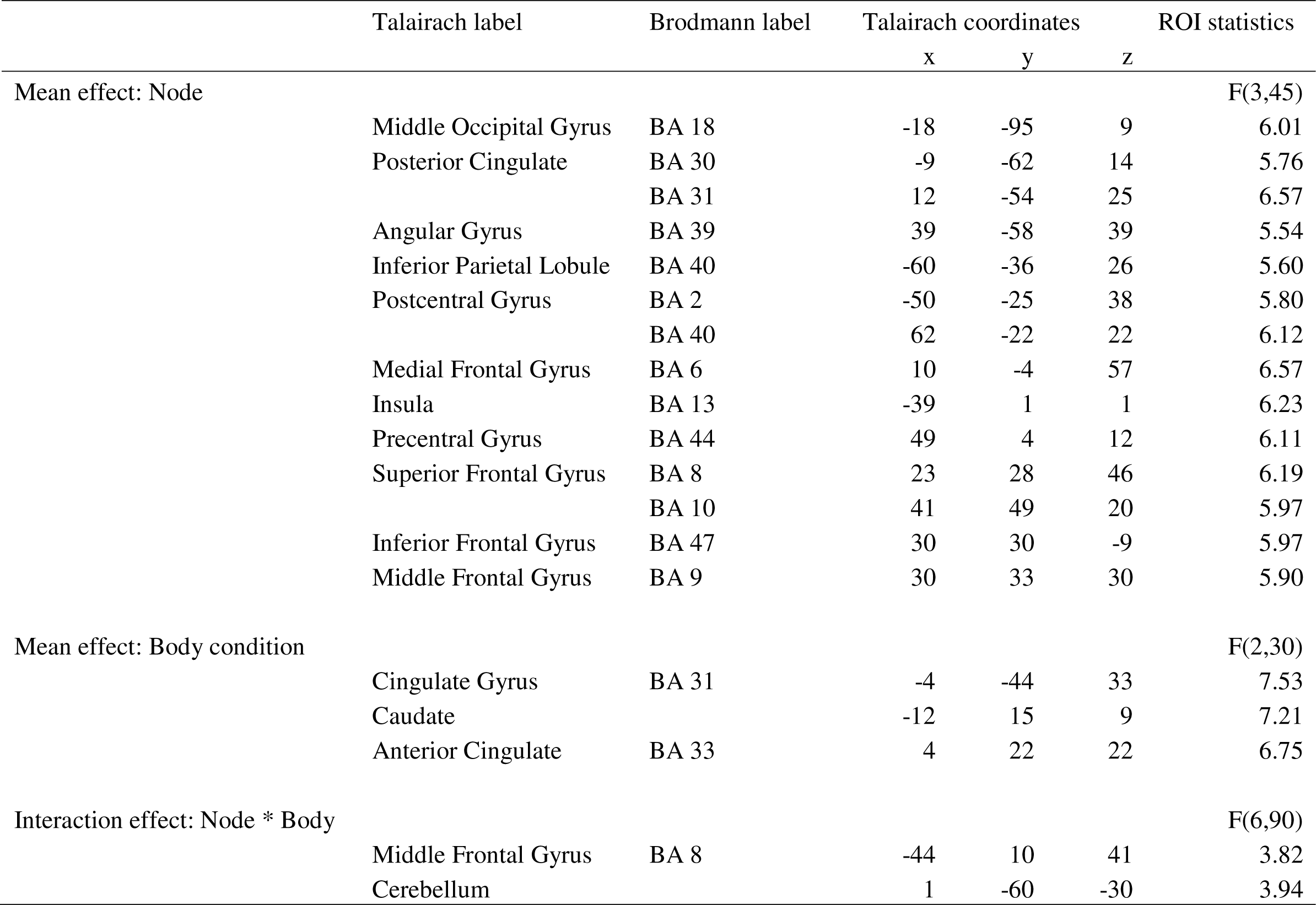
Results from the 3 (body condition) ×4 (node) ANOVA on the connectivity betas.

Within the areas showing a significant main effect of body condition, affective modulations were found in anterior / posterior cingulate cortex (ACC / PCC) and caudate. Subsequent post hoc tests revealed a significant enhancement of overall node connectivity specifically for the defensive body condition (**Figure 5a**; **Table 1**). On the other hand, significant interaction effects between body condition and node node were found in middle frontal gyrus / frontal eye field and in cerebellum. In both clusters, the aggressive body condition significantly increased connectivity to the lpMTG node, while it decreased the connectivity to the LOCs node (**Figure 5b**; **Table 1**).

**Figure 5.**
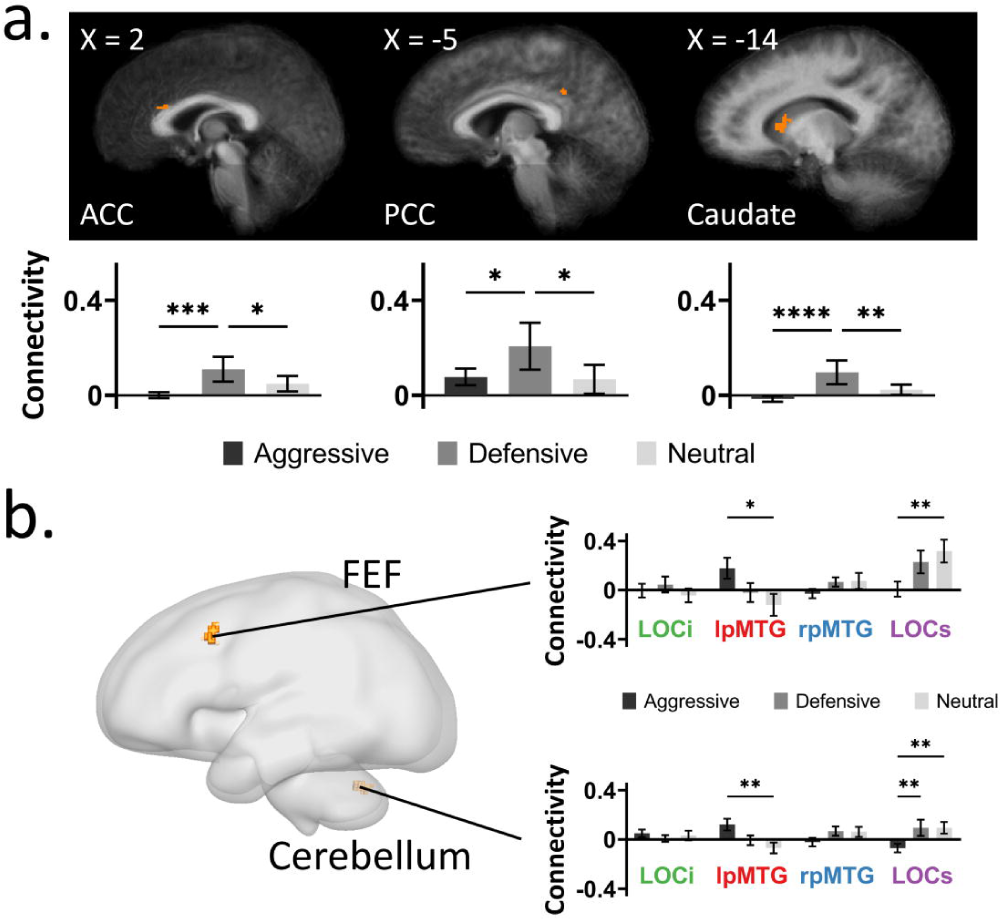
(a) Clusters with significant main effect of the body condition on TOC connectivity (averaged across nodes). (b) Clusters with significant interaction between the body condition and the seed node. Bar plots show the detailed connectivity profile of each cluster. Asterisks indicat the significant pairwise comparisons after Bonferroni correction (*: p < 0.05; **: p < 0.01; ***: p < 0.001; ****: p < 0.0001).

## 4. Discussion

We investigated dynamic whole body processing using data driven methods and a network approach to address three fundamental questions about body and action representation in the brain: (1) is there a network in the brain showing body sensitivity; (2) what is the connectivity profile of the different nodes within this body selective network; and (3) are these nodes differently modulated by action specific information. Our results demonstrated four different nodes involved in body representation, each showing clear anatomical and functional distinctions. Furthermore, each of these nodes had a unique connectivity profile sustaining the processing of different affective whole-body action videos. These data-driven findings on multiple body nodes extend previous proposals based on traditional category localizer methods and provide novel insights into their functional connectivity sustaining action processing.

### 4.1. Four different body selective network nodes

The current study identified four body-selective nodes within the TOC body network (Li et al., 2023). The largest node was the bilateral LOCi located in the inferior division of LOC and the lateral occipital sulcus (LOS). Another node, the bilateral LOCs, was defined in the superior division of LOC. Anterior to the LOCi and LOCs, two unilateral nodes were found in the posterior middle temporal gyrus (MTG), one on the left hemisphere and one on the right (lpMTG and rpMTG). Overall, our findings provide evidence for a distributed network of body representation consistent with an early proposal by Weiner and Grill-Spector (2010). The authors not only identified a single body category, but they also discovered three limb-selective areas organized in a crescent shape around, yet not overlapping with, hMT+. Furthermore, other areas were also involved in limb representation including ITG and MTG. Our results confirm their findings on the anterior-posterior separation of body areas (Weiner & Grill-Spector, 2011). Yet importantly, we extend these findings by demonstrating a distinction between the dorsal and ventral LOC in terms of body processing, challenging the conventional definition of EBA as a homogeneous cluster in previous studies.

To understand differences among the four nodes, we investigated the composition of voxels exhibiting a significantly higher weight for each node compared to the other three. Voxels showing LOCi dominance were found in shape selective areas, such as LO2 and PH (Kolster et al., 2010), as well as in motion and body part selective areas including V4t and FST (Glasser et al., 2016; Kolster et al., 2010). In agreement with our findings, previous studies have suggested that the LOCi region may serve both as an entry of higher-level visual streams as well as a node for integrating the dorsal and ventral stream feedback (Grill-Spector & Malach, 2004; Larsson & Heeger, 2006; Sayres & Grill-Spector, 2008). Recent studies have also found involvement of the LOCi in body kinematics (Marrazzo et al., 2023) and motion feature processing (Robert et al., 2023). Thus, this node may be related to an early integration of the low-/mid-level visual information.

In contrast, LOCs dominance was mainly associated with TOPJ3, a region believed to be modulated by higher-level functions such as theory of mind (ToM) and working memory (Glasser et al., 2016). This once again underscores the functional divergence between the ventral and dorsal LOC. Similarly, both the lpMTG and rpMTG nodes involved multifunctional areas such as PHT and TOPJ2. Both TPOJ3 and PHT/TPOJ2 have shown selectivity for body and face stimuli (Glasser et al., 2016). Interestingly, stronger face form selectivity has been documented for TOPJ3 compared to PHT/TPOJ2, whereas PHT/TPOJ2 has been associated with stronger body form selectivity (Glasser et al., 2016). Thus, the pMTG nodes and the LOCs node may be sensitive to different kinds of visual features associated with specific features of social information.

### 4.2. Different connectivity profiles

An ANOVA with seed nodes and affective conditions as factors was used to compare the nodes’ connectivity profiles. Distinct global connectivity profiles were revealed by the significant main effect of the seed node. The only cluster showing stronger connectivity to the LOCi node was EVA/V2. Regarding to the LOCs node, the strongest connectivity was found with default mode network and dorsal attention network nodes including PCC, precuneus, and superior frontal gyrus (SFG). This is consistent with other studies reporting ventral stream connectivity to the LOC (overlap with the LOCi) and dorsal stream connectivity to the EBA (overlap with the LOCs) (Zimmermann et al., 2018). Compared to the LOCi and LOCs, more widespread connectivity was found for the two anterior nodes, lpMTG and rpMTG, which suggested the pMTG nodes may serve as network hubs connecting the TOC network to the global-level computation. Both pMTG nodes had the highest connectivity to SMG and insula. However, in the case of the rpMTG, this connectivity extended further to encompass the MFG, angular gyrus and SFG. Also, different from the LOCs, the anterior nodes were linked to nodes of ventral attention / salience networks (VAN & SN). The asymmetric results are consistent with the right-literalized distribution of the VAN (Vossel et al., 2014). Thus, these two nodes could contribute to a processing pathway for evaluating both valence as well as relevance in the observer.

### 4.3. Affective modulation for defensive vs. aggressive body movements

The most significant and novel finding derived from our analysis is that for defensive/fearful body images we observed enhanced connectivity between the TOC and the PCC / precuneus as well as the caudate. The former are part of Posteromedial cortex (PMC), known for involvement in episodic memory, including autobiographical memory (Bubb et al., 2017; Leech & Sharp, 2014), and are also part of the default mode network. The PCC has also been shown to exhibit increased activity when attention is directed towards a target of high motivational value (Leech & Sharp, 2014). Furthermore, recent evidence indicates that Anterior Precuneus is causally linked to the bodily self, or the brain resources involved with the observers’ body schema (Lyu et al., 2023). Seen against this background, witnessing fear/defensive actions may engage brain activity intricately associated with the bodily self, in the sense of increasing the involvement of the observer. These results suggest that observing fearful expressions is associated with concomitant neural activity preparing the brain for defensive actions (de Gelder et al., 2004). In support, several brain areas identified in studies of body schema (Berlucchi & Aglioti, 2010), sense of self and its deficits in pathological conditions (Dary et al., 2023) overlap with the network described here.

In contrast, the connectivity modulation for aggressive actions showed a different pattern and varied across different nodes. As revealed by the ANOVA analysis, a significant interaction effect was observed for FEF and cerebellum regions. When viewing the aggressive videos, the two regions showed increased connectivity to the lpMTG node, and decreased connectivity to the LOCs node. As mentioned above, both lpMTG and LOCs were composed of voxels from social perception regions such as TPOJ and PHT and may be related to the processing of different types of social features. And compared to LOCs, the lpMTG node exhibited a higher level of global connectivity. Moreover, the FEF region is known for its involvement in shaping attention (Veniero et al., 2021) and as such the FEF may be part of decision making (Krajbich et al., 2021). Thus, the opposite FEF connectivity pattern to the lpMTG and LOCs may suggest the reorientation of salient features and the distributed computation during the presence of aggressive stimuli.

## 5. Conclusions

We showed that body representation is implemented in the brain in several different body sensitive hubs that each have their connectivity network. These hubs and connectivity networks have a relatively specific functional role in body and action perception. This supports the notion that these are body sensitive hubs that are not so much defined by abstract body category selectivity than by their different functional roles in action understanding.

## Acknowledgments

This work was supported by the European Research Council ERC Synergy grant (Grant agreement 856495, Relevance), by the European Union’s Horizon 2020 research and innovation programme (Grant agreement 101017884, GuestXR), the European Union’s Horizon Europe research and innovation programme (Grant agreement 101070278, ReSilence), and by the Future and Emerging Technologies (FET) Proactive Program H2020-EU.1.2.2 (Grant agreement 824160, EnTimeMent).

